# Impact of vertebrate host parasitaemia on *Plasmodium* development within mosquitoes

**DOI:** 10.1101/2024.07.22.604581

**Authors:** Julie Isaïa, Molly Baur, Jérôme Wassef, Sarah Monod, Olivier Glaizot, Philippe Christe, Romain Pigeault

## Abstract

**Background:** In vector-borne diseases, invertebrate hosts are exposed to highly variable quantities of parasites during their blood meal. This heterogeneity may partly explain the overdispersed distribution of parasites within the vector population, as well as the variability in the extrinsic incubation period (EIP) of the parasite. Indeed, the quantity of parasites ingested is often considered as a good predictor of the quantity of parasites that will develop within the vectors, as well as the speed at which they will develop (EIP). However, density-dependent processes can strongly influence the relationship between parasite burden in the vertebrate host and in vectors, making this relationship not always clear.

**Methods:** Here, we used the avian malaria system to investigate whether the proportion of red blood cells infected by sexual and/or asexual stages of malaria parasite influences the intensity of malaria infection and the EIP of *Plasmodium* within the invertebrate vectors. For this purpose, we have experimentally infected twelve vertebrate hosts in order to generate a range of intensity of infection. More than a thousand mosquitoes took a blood meal on these hosts and the development of *Plasmodium* within the vectors was followed for more than 20 days.

**Results:** The main finding presented in this study reveals a negative relationship between the intensity of infection in the vertebrate host and the EIP. Four days were sufficient for 10% of infected mosquitoes fed on the most infected hosts to become infectious. However, the number of transmissible stages did not significantly vary according to the vertebrate host intensity of infection.

**Conclusion:** While the quantity of ingested parasites had no impact on the density of transmissible stages in infectious mosquitoes, the EIP was affected. Studies have demonstrated that small changes in the EIP can have a significant effect on the number of mosquitoes living long enough to transmit parasites. Here, we observed a difference of 4-6 days in the detection of the first transmissible stages, depending on the intensity of infection of the bitten vertebrate host. Considering that a gonotrophic cycle lasts 3-4 days, the shortened EIP observed here may have significant effects on *Plasmodium* transmission.

## Introduction

The rate at which a population grows or declines is influenced by environmental conditions but also by the number of individuals already present in the population. This phenomenon, studied in population ecology, is known as density-dependence processes [1–4]. Density-dependent mechanisms are considered of central importance in stabilizing the population growth of both free-living [5] and parasitic organisms [6,7]. Parasites are especially subject to density-dependent processes as they often have complex life cycles involving bottlenecks upon transmission followed by exponential growth [2,7–9]. Positive but also negative density-dependent processes are common in host-parasite associations and both may influence parasite transmission rate [10,11], resilience of an infection and control interventions [12,13]. Since parasites are typically overdispersed within host populations (i.e., are aggregated and the variance is considerably greater than the average [14,15]), the severity of density-dependent processes can differ greatly among individuals and consequently, their effects at the population level can be strongly influenced by the degree of parasite aggregation [7,16]. Understanding the causes of parasite overdispersion is therefore an important step towards assessing the impact of density-dependent processes on parasite transmission dynamics [17].

Parasite overdispersion has been widely described in several vector-borne diseases [18–21]. However, most studies have focused on the causes and consequences of aggregate parasite distribution in vertebrate host populations, leaving aside what happens within invertebrate host populations (i.e. the vector). Yet, several key features that determine parasite transmission dynamics in vertebrate host populations (e.g., individual vectorial capacity, [22]) are highly dependent on the intensity of parasite infection in the vector [23–26]. Fluctuations in parasite density within vector population may be due to the genetic background or age of the invertebrate hosts, which influence their susceptibility to parasite infection [18,27,28]. Environmental factors may also be involved, such as temperature which affects the parasite’s ability to complete its life cycle within the invertebrate host [29,30]. The amount of parasite ingested by the vector during their infestation (usually a blood meal) can also generate infection heterogeneity (e.g., [31,32]). Indeed, parasite numbers vary between vertebrate hosts and also spatiotemporally within individuals [33,34]. Although a positive correlation is expected between the quantity of parasite present in the vertebrate host blood and the density of transmissible stages of the parasite found in vectors (after the incubation period), the relationship is not always obvious. This is particularly true for *Plasmodium* parasites, the aetiological agents of one of the most studied vector-borne diseases, malaria. The relationship between the proportion of red blood cells of the vertebrate host infected by gametocyte (i.e., the transmissible stage) and the parasite burden in the mosquito seems to be specific to *Plasmodium-*mosquito associations. For the human malaria parasite *P. falciparum* and both *An. coluzzii* and *An. gambiae* mosquitoes, the parasite burden within mosquito midgut (i.e. the oocyst stage) increases with the gametocytes density [35–38]. On the other hand, this relationship is either negative or unclear in the case of rodent and avian malaria, respectively [39,40].

To date, the majority of the studies trying to elucidate the relationship between vertebrate host gametocytaemia (i.e., percentage of red blood cells infected by a mature gametocyte) and the parasite burden in mosquitoes only consider the oocyst stage, which is the simplest and least expensive measure of the mosquito infection. Even though oocyst burden seems to be positively correlated with the likelihood of transmitting sporozoites to a new vertebrate host (Miura *et al.* 2019) the extrinsic incubation period of the parasite (EIP) should be considered [43]. EIP refers to the time required for the parasite to replicate and disseminate to the mosquito salivary glands after the ingestion of the infected blood meal. It determines the time at which the vector will be infectious and able to transmit the parasite to a new vertebrate host; the shorter the EIP the greater the opportunity for the mosquito to transmit the parasite.

The aim of this study was to assess the role of the vertebrate host parasitaemia (i.e., percentage of red blood cells infected by either sexual or asexual stages of *Plasmodium*) and gametocytaemia on the within-vector parasite infection dynamics. We investigated whether the parasitaemia and gametocytaemia in the vertebrate host may determine the density of oocysts in mosquito midgut and/or the extrinsic incubation period of *Plasmodium*. We predict that mosquitoes biting highly infected hosts will have higher oocyst burdens and be infectious earlier than mosquitoes feeding on hosts with low parasitaemia and gametocytaemia. To test this prediction, we used the natural avian malaria system [40], with domestic canaries (*Serinus canaria*) as vertebrate host, an haemosporidian parasite recently isolated from the wild (*Plasmodium relictum*) and a natural mosquito vector in the field (*Culex pipiens* [44]).

## Materials and Methods

### Biological material

The experiment was carried out using a *Plasmodium relictum* (lineage SGS1) strain isolated from an infected house sparrow (*Passer domesticus)* captured in December 2020 on the campus of the University of Lausanne, Switzerland (46°31’25.607’’N 6°34’40.714’’E). This strain was maintained through regular passages across our stock canaries (*Serinus canaria*) using intraperitoneal (i.p.) injections until the beginning of the experiment [40].

The *Culex pipiens* mosquito population used in this experiment was initiated from wild clutches collected in Lausanne in August 2017 and maintained in the insectary since. Mosquitoes were reared using standard protocols [45]. On the day prior to mosquito blood meal, around 1000 7-10-day-old females were haphazardly captured from different emergence cages and placed inside experimental cages (90 females per cage, L40 x W40 x H40 cm). During this time, females were deprived of sugar solution, but had access to water, to maximize the biting rate.

### Experimental design

#### Infection of vertebrate hosts

Twelve domestic canaries (*Serinus canaria*) were infected with *P. relictum* (lineage SGS1) by i.p. from our stock canaries. One bird died day 5 post-infection and was therefore removed from the analyses. The dynamic of infection was monitored for each bird every two days from day 5 to day 20 post-infection by measuring the parasitaemia and the gametocytaemia by microscopic examination using Giemsa-stained blood-smear. Blood samples (3-5µL) were taken from the medial metatarsal vein. The number of red blood cells infected by sexual and/or asexual *Plasmodium* stages were counted per 3000-4000 erythrocytes in randomly chosen fields on the blood smears [46].

#### Mosquito exposition to P. relictum

On day 12 post vertebrate host infection, corresponding to the peak of infection for our parasite strain, birds were exposed individually to 90 uninfected mated female mosquitoes for 3h (6-9 p.m.). At the end of the experiment, blood-fed females were counted and kept individually in plastic tubes under standard laboratory conditions (25°C - 70%RH) with 10% *ad libitum* glucose.

#### Mosquito dissections

Every two days starting from day 4 to day 20 post-blood meal, 4 ± 1 mosquitoes per bird were haphazardly sampled in order to monitor both the dynamic of oocysts formation and sporozoites production. Each mosquito was dissected to (i) count the number of oocysts in their midgut with the aid of a binocular microscope and (ii) quantify the transmissible sporozoites in their head/thorax using real-time quantitative PCR (see molecular analyses below). The appearance of sporozoites in the head/thorax homogenate is strongly correlated with the appearance of sporozoites in the salivary glands [49]. The amount of haematin deposited in the individual resting tubes was used to estimate the size of the blood meal (see [45]).

#### Molecular analyses

Real-time quantitative PCR was used to study the presence/absence and the density of *Plasmodium* sporozoites in the mosquito head/thorax. Beads were added to each head/thorax samples and flash frozen in liquid nitrogen in order to grind the samples. DNA was then extracted using the Qiagen DNeasy Blood and Tissue Kit following the manufacturer’s instructions, except that the samples were incubated overnight. For each individual (i.e., mosquito), two qPCRs were carried out: one targeting the mtDNA cytb gene of *Plasmodium* (Primers L4050Plasmo 5’-GCTTTATGTATTGTATTTATAC-3’, H4121Plasmo 5’-GACTTAAAAGATTTGGATAG-3’, Probe TexasRed-CYTB-BHQ2 5’-CCTTTAGGGTATGATACAGC-3’) and the other targeting the CQ11 gene of *Cx. pipiens* mosquitoes (Primers 1725-F 5’-GCGGCCAAATATTGAGACTT-3’, 1726-R 5’-CGTCCTCAAACATCCAGACA-3’, Probe FAM-CQ11-BHQ1 5’-GGAACATGTTGAGCTTCGGK-3’). All samples were run in triplicate (QuantStudio 6 and 7 Pro Real-Time PCR Systems) and samples with a threshold Ct value higher than 35 for the parasite were considered uninfected. Relative quantification values (RQ) were calculated to assess the parasite prevalence and can be interpreted as the fold-amount of target gene (*Plasmodium* CYTB) with respect to the amount of the reference gene (*Cx. pipiens* CQ11) and are calculated as 2^-(C*t*CYTB *Plasmodium* - C*t*CQ11 *Cx. pipiens*)^.

#### Statistical analyses

Analyses were carried out using the R statistical software (v. 4.2.1). The relationship between birds’ gametocytaemia and parasitaemia was studied using a linear regression. We verified the model’s assumptions by plotting residuals using the simulateResiduals function from the DHARMa package [50]. This checking procedure was applied to all the models presented below. Because of the high degree of collinearity between parasitaemia and gametocytaemia, these two variables were never fitted as response variables in the same model. The influence of parasitaemia or gametocytaemia on the day post-blood meal at which the maximum burden of oocysts or sporozoites was reached as well as the day at which 10% of the mosquitoes in a batch were detected as sporozoite carriers (i.e., EIP10) was analyzed using linear models. Influence of parasitaemia or gametocytaemia on mosquitoes’ blood meal size, oocyst burden and sporozoites density were analyzed fitting bird as a random factor into the models using lmer or glmer.nb (package: lme4, [51]) according to whether the errors were normally (haematin quantity, sporozoites density) or negative binomially distributed (oocyst burden). Where necessary, blood meal size and oocyst burden were added as response variables in the models. Maximal models, including all higher-order interactions, were simplified by sequentially eliminating non-significant terms and interactions to establish a minimal model [52]. The significance of the explanatory variables was established using a likelihood ratio test which is approximately distributed as a Chi-square distribution or a F test [53]. The significant Chi-square or F values given in the text are for the minimal model, whereas non-significant values correspond to those obtained before the deletion of the variable from the model. The different statistical models built to analyse the data are described in electronic supplementary material in Table S1.

## Results

### Parasitaemia and gametocytaemia of vertebrate host

This experiment was carried out to investigate whether parasitaemia and gametocytaemia in the vertebrate host the day of the infectious blood meal underpinned the parasite development within the invertebrate vectors. Eleven vertebrate hosts out of the twelve experimentally infected survived until day twelve, allowing a range of parasitaemia and gametocytaemia values to be generated. The parasitaemia on the day of the mosquito blood meal (i.e., day 12 post-infection) followed an aggregated distribution (variance to mean ratio = 6.89, mean ± sd: 3.47 ± 4.89). Three hosts had a parasitaemia of less than 1%, four individuals had a parasitaemia ranging between 1 and 2% and four individuals had a parasitaemia of over 3%, one of which presented a parasitaemia of over 17% (**Fig. 1A**). The percentage of red blood cells infected with mature gametocytes was on average low (mean ± sd: 0.0015 ± 0.002) but, as reported in a previous study (e.g., [40,54]), we observed a strong positive correlation between parasitaemia and gametocytaemia (model 1: F = 27.68, p < 0.001, Adjusted R-squared: 0.724, **Fig 1A**). As the conclusions of the subsequent analyses are essentially the same whether parasitaemia or gametocytaemia was fitted as explanatory variable, we focus hereafter solely on the influence of parasitaemia. Parasitaemia measurement was indeed much less prone to slight error, since it only involved counting the number of infected red blood cells. In the case of gametocytaemia measurement, it was necessary to differentiate the parasite stages and their maturation levels, which can sometimes be a little subjective (at present, there is no molecular tools for identifying mature gametocytes). All analyses incorporating gametocytaemia instead of parasitaemia fitted as explanatory variable are however presented in **Appendix 1**.

**Fig. 1.**
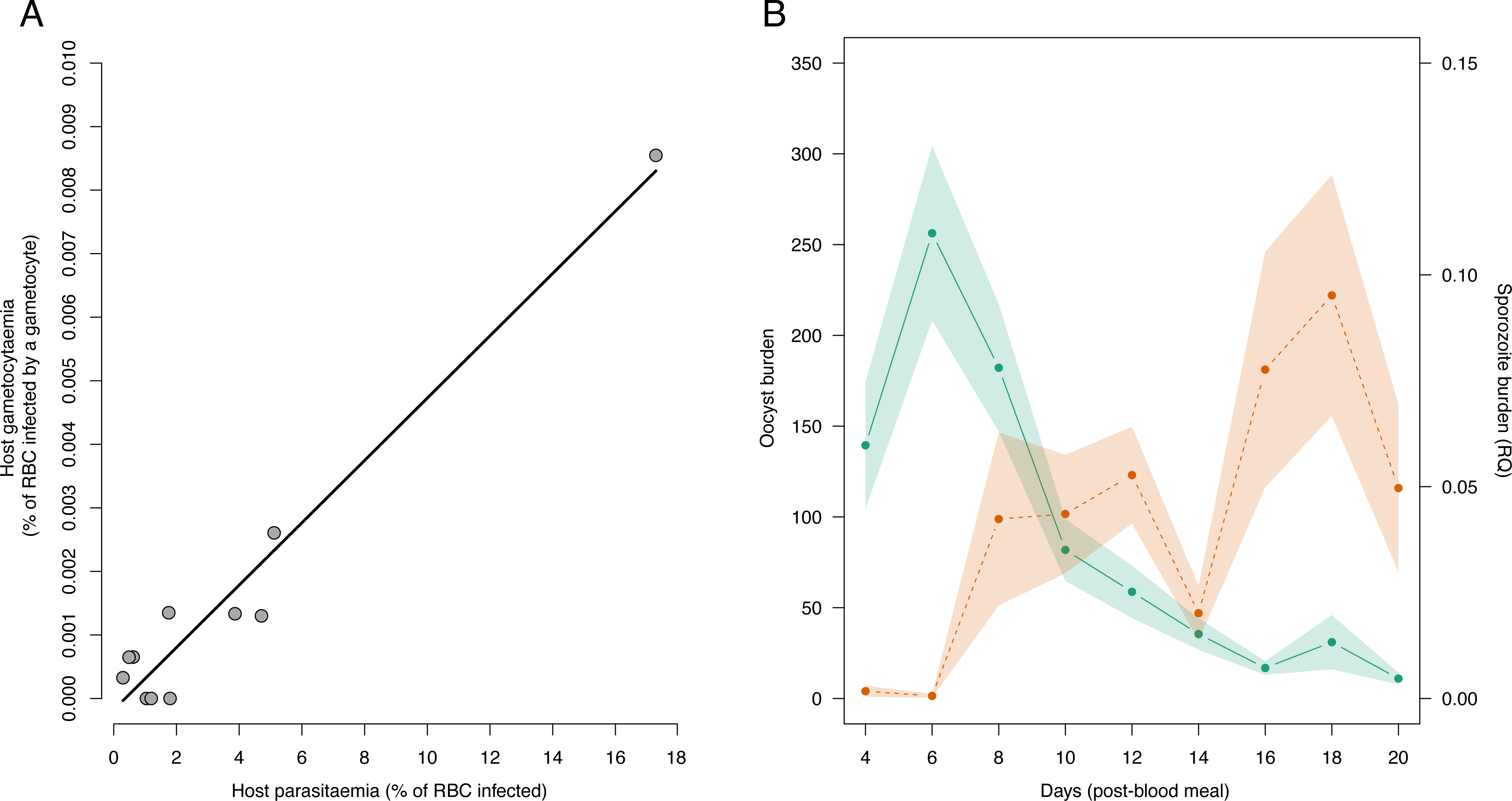
(**A**) **Relationship between vertebrate host parasitaemia and gametocytaemia**. Eleven individuals were included in this analysis. (**B**) **Temporal dynamics of *Plasmodium* development in mosquitoes**. Average oocyst burden (green, left axis) and sporozoite counts (salmon, right axis) at each dissection day. Green and salmon shadows represent standard error. The left axis represents the average number of oocysts counted per female. The right axis represents the amount of sporozoites quantified by qPCR.

### Temporal dynamics of *Plasmodium* development in mosquitoes

The temporal dynamics of oocyst production inside the mosquito midgut were consistent between mosquito batches fed on the eleven infected birds (**Fig. 1B**, Fig. S1). Oocysts were detected in all batches of mosquitoes from the first day of dissection, i.e. the fourth day after the blood meal. The peak oocyst burden was reached between days 6 and 8 (**Fig. 1B**, Fig. S1). Subsequently, a swift reduction in the oocyst count was noted, such that by day 14 post-blood meal, only a residual burden ranging from 1 to 20% of the peak oocysts persisted. The first sporozoites appear on the head-thorax homogenate mainly after day 6 (i.e. EIP, **Fig. 1B**).

However, for two mosquito batches sporozoites were detected in some females on day 4 post-blood meal (**Fig. S1**). The peaks concentration of sporozoites were reached between day 10 and 12 post-blood meal for half of the batches of mosquitoes and between day 16 and 18 for the other half (**Fig. 1B**, Fig. S1).

### Effect of vertebrate host parasitaemia on oocyst production

The size of the blood meal of mosquitoes was not influenced by the parasitaemia of the bird on which they fed (mean ± se, 20.07µg ± 0.055, model 2: X^2^=0.256, p = 0.613). Since all mosquitoes already had oocysts in their midgut when we started the dissections (day 4 post-blood meal), we were unable to study the relationship between parasitaemia and the first appearance of oocysts. We showed that parasitaemia of vertebrate hosts had no influence on the day post-blood meal on which the oocyst peak was reached (model 3: X^2^=1.29, p = 0.25, **Fig. 2A**) or on the density of oocysts reached during this peak (model 4: X^2^=2.334, p = 0.127, **Fig. 2B**).

**Fig. 2.**
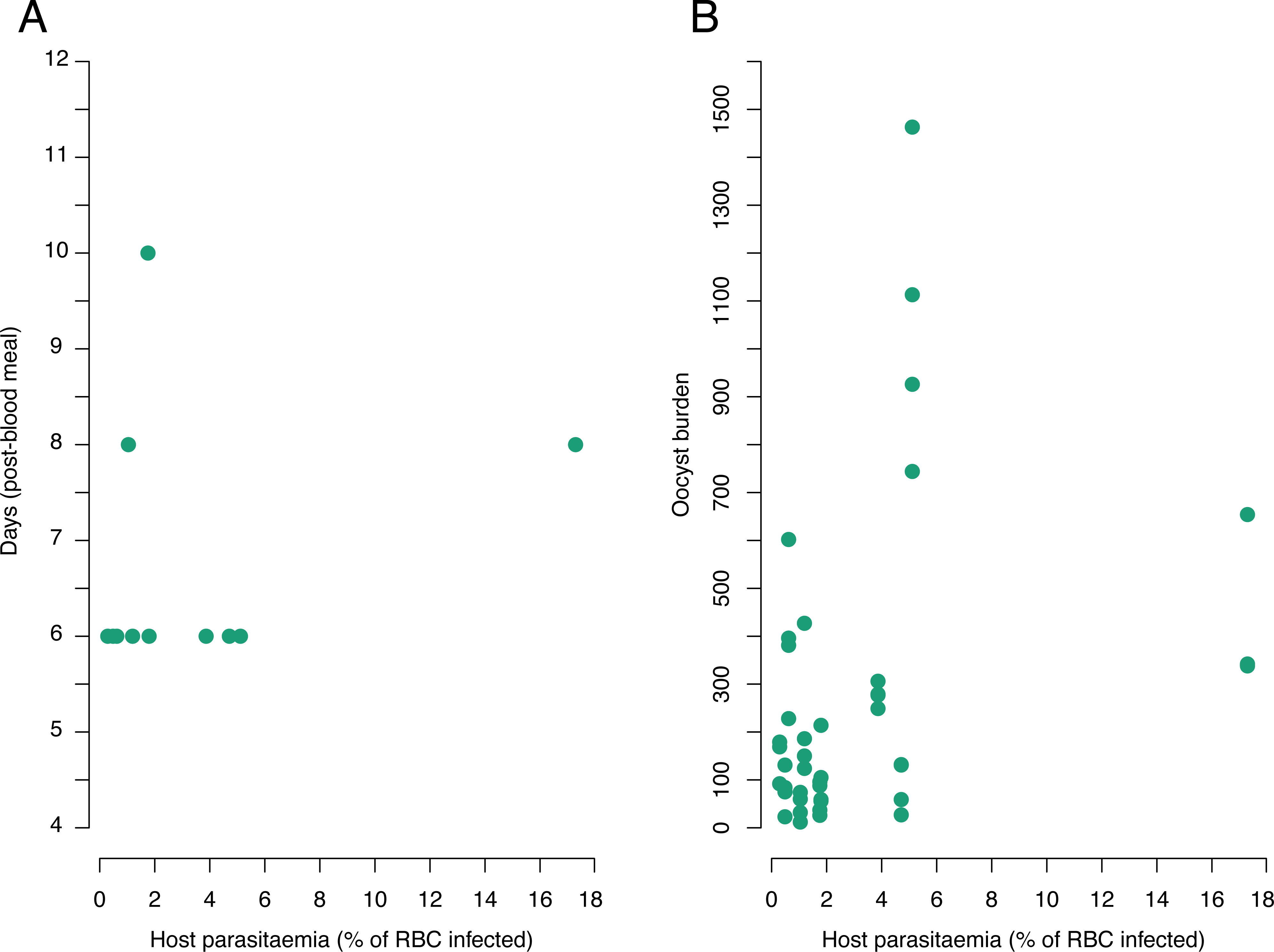
Relationship between vertebrate host parasitaemia and development of oocysts in the midgut of mosquitoes. (**A**) Influence of parasitaemia on the day post-blood meal at which the maximum oocyst burden was reached. Each green dot corresponds to the batch of mosquitoes fed on a vertebrate host where the maximum oocyst burden was reached. (**B**) Influence of parasitaemia on oocyst burden measured on the day when oocyst burden was maximum within a focal mosquito batch. Each dot represents the number of oocysts in the midgut of a mosquito.

### Effect of vertebrate host parasitaemia on sporozoite production

The time required for sporozoites to be detected in the head-thorax homogenate of 10% of infected mosquitoes was influenced by the parasitaemia of the vertebrate hosts on which the mosquitoes fed (EIP10, model 5: F = 8.29, p = 0.020, **Fig. 3A**). Batches of mosquitoes that had bitten the most infected vertebrate hosts showed sporozoites as early as day 4 post-blood meal, which was up to four to six days earlier than females that had bitten the least infected vertebrate hosts (**Fig. 3A)**. Fitting the quadratic term (parasitaemia²) slightly improved the model fit (model 5: X^2^ = 5.036, p < 0.0493), suggesting the day at which sporozoites were detected was a decelerating polynomial function of vertebrate host parasitaemia. As previously reported, peak sporozoite concentration in mosquitoes were reached between day 10 and 18 post-blood meal, but the delay to reach it was not influenced by vertebrate host parasitaemia (model 6: F = 0.917, p = 0.363). The maximum sporozoite density reached during the peak was not influenced by the parasitaemia of the vertebrate hosts either (model 7: X² = 0.054, p = 0.815) but was influenced by the oocyst load reached during the oocyst peak (model 7: X² = 7.575, p = 0.006; **Fig. 3B**). The higher the oocyst burden, the higher the sporozoite load.

**Fig. 3.**
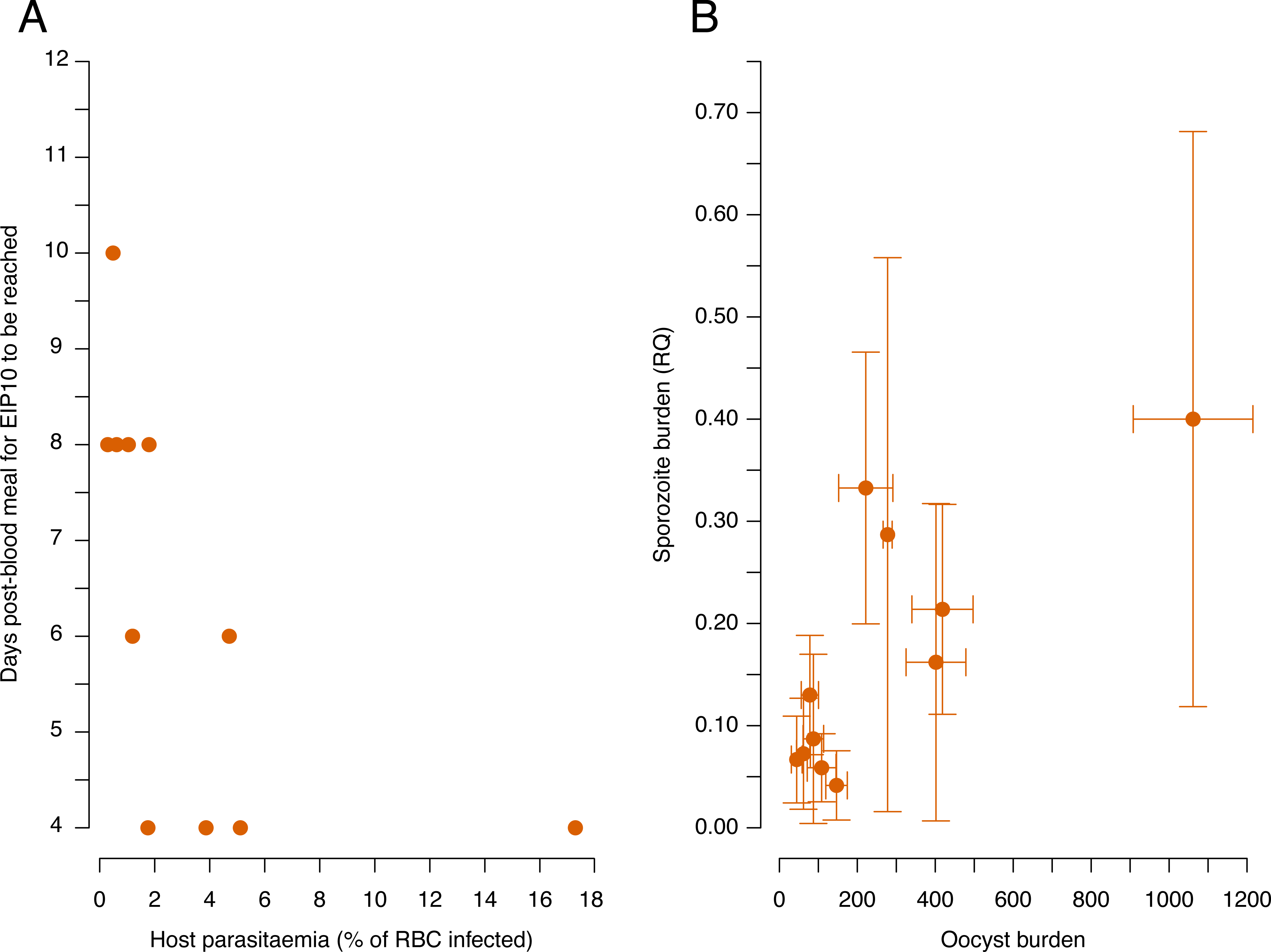
(**A**) **Influence of parasitaemia on the day post-blood meal at which the EIP10 was reached** (i.e., at least ten percent of mosquitoes carry sporozoites). Each salmon dot corresponds to a batch of mosquitoes fed on a vertebrate host. (**B**) **Relationship between average oocyst burden and average sporozoite load quantified** in each batch of mosquitoes on the day of maximum oocyst and sporozoite load respectively.

## Discussion

The main finding presented in this study reveals a negative relationship between the intensity of malaria infection in the vertebrate host and the delay required after biting for the mosquito vectors to become infectious. We showed that the time required for 10% of infected mosquitoes to become infectious (EIP10) was reached by the batches of vectors fed on the most infected vertebrate hosts as early as day four after the blood meal. However, taking a blood meal on highly infected vertebrate hosts did not affect the intensity of parasite infection within the mosquito: the number of oocysts, as well as the concentration of sporozoites, is not significantly affected by the vertebrate host parasitaemia or gametocytaemia.

In several natural systems such as *P. falciparum* and *An. coluzzii*, *P. falciparum* and *An. gambiae, P. vivax* and *An. dirus* or *P. mexicanum* and *Lutzomyia vexator,* the oocyst burden in the invertebrate vector is positively correlated to the gametocyte densities [31,38,39,55]. This relationship, however, was not very strong in many studies and there was considerable variability in the number of oocysts associated with a given parasitaemia or gametocytaemia value (e.g., [55–58]). Density-dependent processes are at play in the regulation of the parasite journey within the vector with major bottlenecks occurring during life-stages transition from gametocytes to oocysts [9,59–61]. Here, we observe that the maximum oocyst burden reached during peak oocyst development was not significantly influenced by parasitaemia or gametocytaemia, although we reported a positive trend between gametocyte burden and oocyst burden. The number of vertebrate hosts included in the study and the widely dispersed distribution of gametocytaemia (the majority of birds have very low gametocytaemia levels) could explain why the relationship between these two parameters was not statistically significant. Similar conclusions were however reported in a previous study carried out with the same biological system on forty birds with a much wider range of gametocytaemia (i.e., 0.1 to 9%, [40]). Therefore, our results and those published previously point in the same direction and strongly suggest that in the natural *P. relictum* / *Cx. pipiens* system, oocyst burden is not strongly influenced by gametocyte density [33,40]. This could be explained by the fact that not all gametocytes counted in blood smears were yet infectious (i.e., ready to produce micro-or macro-gametes in the mosquito’s midgut). The mechanisms underlying gametocyte infectivity remain poorly understood. Although we know that gametocytes go through several stages of development before reaching the mature stage [62], we do not know whether the mature stage is systematically infectious.

Parasitaemia and gametocytaemia in birds also had no influence on the maximum sporozoite density measured in the head-thoraxes of *Cx. pipiens.* However, we observed a positive relationship between maximum sporozoite density and the maximum average oocyst burden. Although the number of sporozoites produced per oocyst can vary from a few dozen to several thousand depending on the parasite/mosquito pair [63], a positive correlation between these two parasites stages is commonly reported [49,55,64–66]. The strength of the relationship between oocyst burden, sporozoite density and the rate of transmission of *Plasmodium* to vertebrate hosts is however difficult to assess due to the methodological complexity associated with the quantitative monitoring of the different parasite stages within a single mosquito. Nonetheless, a recent study conducted on both human and rodent malaria systems has clearly demonstrated a significant positive correlation between mosquito parasite load (oocyst and sporozoite stages) and the quantity of sporozoites injected during probing (i.e., inoculum size, [66]).

In addition to inoculum size, the time lapse between mosquito infection (i.e., ingestion of gametocytes during the blood meal) and the moment when they become infectious (i.e., EIP) also has a major influence on the transmission dynamics of *Plasmodium*. Small changes in EIP can have a large effect on the number of mosquitoes living long enough to transmit parasites and subsequently on the number of hosts a vector might infect over its lifespan (e.g., [28,67]). For the majority of *Plasmodium* species, EIP is on average between 8 and 16 days (see Table 1 in [68] as well as e.g., [69,70] for studies carried out on *P. relictum*). Several studies have shown that EIP can be highly heterogeneous between mosquitoes due to differences in nutrition (e.g., [48,67,71,72]) and environmental conditions (e.g., temperature, see figure 1 in [43]), however, considerable variation remains even after accounting for these factors. Our results suggest that variation in EIP could be explained in part by the quantity of parasites ingested by the mosquitoes. Indeed, we detected the first sporozoites in batches of mosquitoes fed on the most infected hosts, on average four to six days earlier than in batches of mosquitoes fed on hosts characterized by lower parasitaemia and gametocytaemia. Interestingly, a previous study carried out on the same biological system, but with only three infected vertebrate hosts, highlighted a similar trend (see Fig. S1 in [73]). Considering that *Cx. pipiens* gonotrophic cycle lasts 3 to 4 days, shortening the EIP in response to over-crowding, when vector’s resources become limited, might result in a form of fertility insurance for the parasite. Besides the potential to infect more hosts overall, faster development may be adaptive when there is a risk that the vector will not live long enough to transmit the parasite.

This pattern may be explained by density-dependent processes which may operate during the first steps of *Plasmodium* development within mosquito midgut [9,24,74]. A high gametocyte density in the blood meal may lead to competition between parasites at different step of early sporogony (e.g., fecundation, resources, colonization of the midgut wall). This competition could either accelerate the development of all parasites and/or favor fast-growing parasite variants, both leading to early development of the first oocysts and, consequently, the rapid appearance of the first sporozoites. A theoretical study supports this hypothesis, but their results also suggest that above a certain density, the number of parasites in the mosquito midgut is so high that it could have a negative effect on the EIP (e.g., by negatively impacting oocyst survival rates or delaying oocyst burst, for example, [75]).

## Conclusions

In summary, our study has demonstrated for the first time that the intensity of infection within the vertebrate host on which a mosquito feeds influences the extrinsic incubation period of *Plasmodium*. The first sporozoites were indeed observed in the batches of mosquitoes fed on the most infected hosts. It should be noted, however, that in this study we only used acutely infected hosts, and therefore with high parasitaemia. Indeed, as this was a pioneering study, we wanted to generate the widest possible range of infection intensities to detect a signal that might have been weak. In the field, the vast majority of vertebrate hosts are in chronic stage of infection [76–78] and it would then be relevant to determine whether the results we obtained are transposable to milder parasitaemia.

## Supplementary information

Additional file 1. Fig. S1.

Additional file 2. Table S1.

## Abbreviations

EIP: Extrinsic incubation period

RH: Relative humidity

## Declarations

### Author’s contributions

J.I., R.P., O.G. and P.C. conceived the study and elaborated the experimental design. J.I., M.B. and S.M. performed the experiments. M.B. and J.W. did the molecular analyses. J.I. and R.P. analyzed the data. J.I. wrote the first draft of the manuscript and all authors contributed substantially to revision.

### Ethics approval and consent to participate

This study was approved by the Ethical Committee of the Vaud Canton veterinary authority, authorization number 1730.4.

## Consent for publication

All participants consented to have their data published.

