## Supplemental information for "Impact of vertebrate host parasitaemia on *Plasmodium* development within mosquitoes"

**Fig. S1 Temporal dynamics of *Plasmodium* development in batches of mosquitoes for each bird bitten/parasitaemia**. Average oocyst burden (green, left axis) and sporozoite counts (salmon, right axis) in mosquitoes for each bird (parasitaemia) at each dissection day. Green and salmon shadows represent standard error. The left axis represents the average number of oocysts counted per female. The right axis represents the amount of sporozoites quantified by qPCR.

**Table. S1 Description of statistical models used in the study.** “N” gives the sample size. “Maximal Model” includes the complete set of explanatory variables. “Minimal model” gives the model containing only the significant variables and their interactions. Square brackets indicate variables fitted as random factors. Curly brackets indicate the error structure used (n: normal errors, b: binomial errors). The response variable was not transformed unless otherwise stated.

Fig. S1

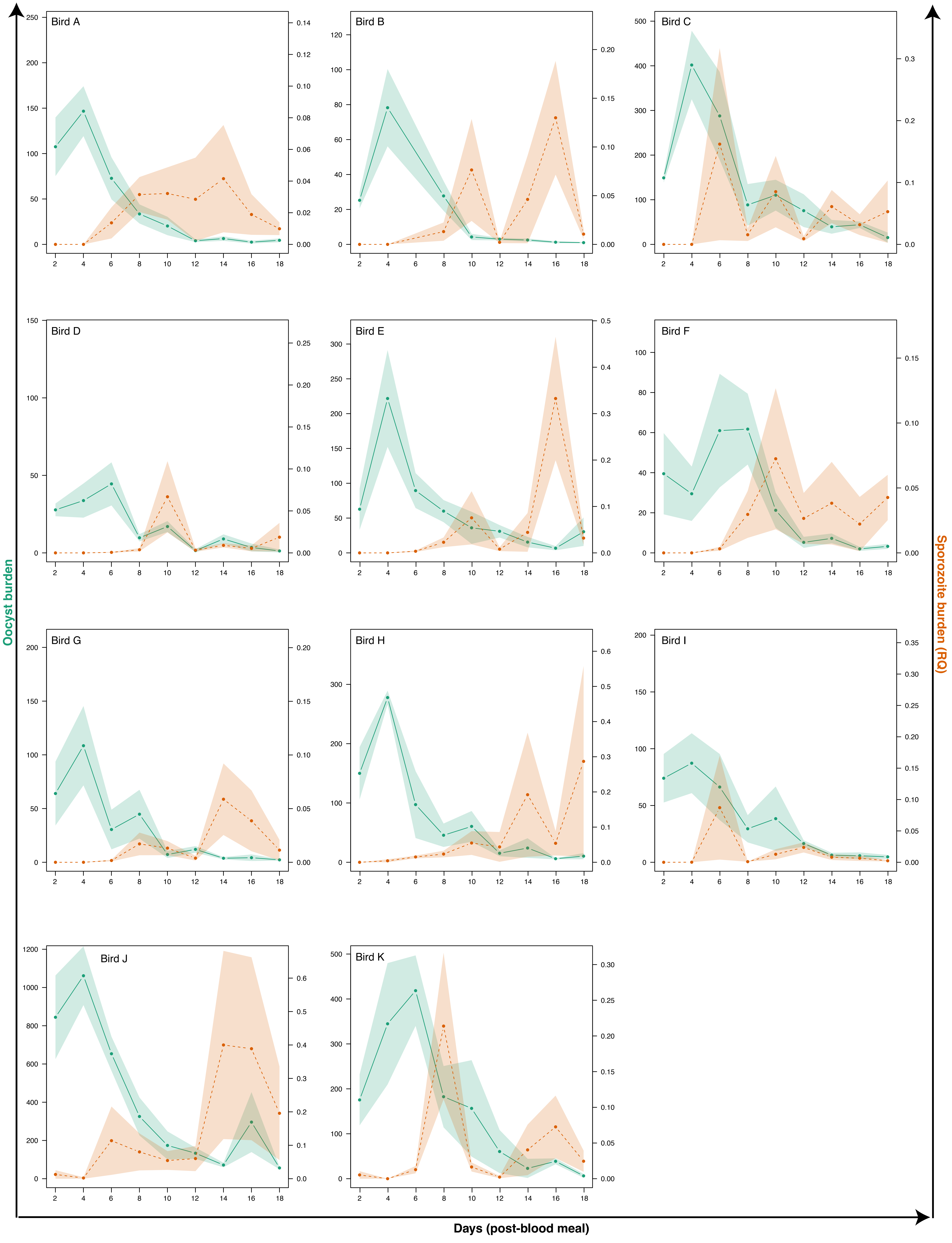
Bird A
Bird B
Bird C
Bird D
Bird E
Bird F
Bird G
Bird H
Bird I
Bird J
Bird K
Oocyst burden
Sporozoite burden (RQ)
Days (post-blood meal)

### Table S1

| Variable of interest | Resp. variable | Model nb | N | Maximal model | Minimal model | R subroutine {error struct} |
|---|---|---|---|---|---|---|
| Gametocytaemia on the day of the mosquito blood meal | sqrt(Gam) | 1 | 11 | Parasitaemia | Parasitaemia | lm {n} |
| Blood meal size | Haematin | 2 | 403 | Parasitaemia + (1\|Bird) | 1 + (1\|Bird) | lmer {n} |
| Day post-blood meal on which the oocyst peak was reached | Day | 3 | 11 | Parasitaemia | 1 | lm {n} |
| Oocyst burden | Oocyst | 4 | 46 | Parasitaemia + Haematin + (1\|Bird) | Haematin + (1\|Bird) | glmer.nb |
| Time required for sporozoites to be detected in 10% of infected mosquitoes | EIP10 | 5 | 11 | Parasitaemia + Parasitaemia^2 | Parasitaemia + Parasitaemia^2 | lm {n} |
| Day post-blood meal on which the sporozoite peak was reached | Day | 6 | 11 | Parasitaemia | 1 | lm {n} |
| Sporozoite burden | Sporo | 7 | 41 | Parasitaemia + Haematin + Oocyst + (1\|Bird) | Oocyst + (1\|Bird) | lmer {n} |
